# Pharmacogenomics, biomarker network and allele frequencies in colorectal cancer

**DOI:** 10.1101/561316

**Authors:** Andrés López-Cortés, César Paz-y-Miño, Santiago Guerrero, Gabriela Jaramillo-Koupermann, Dámaris P. Intriago-Baldeón, Jennyfer M. García-Cárdenas, Patricia Guevara-Ramírez, Isaac Armendáriz-Castillo, Paola E. Leone, Luis Abel Quiñones, Juan Pablo Cayún, Néstor W. Soria

## Abstract

Colorectal cancer (CRC) is one of the leading causes of cancer death worldwide. Over the last decades, several studies have shown that tumor-related genomic alterations predict tumor prognosis, drug response and toxicity. These observations have led to the development of a number of precision therapies based on individual genomic profiles. As part of these approaches, pharmacogenomics analyses genomic alterations that may predict an efficient therapeutic response. Studying these mutations as biomarkers for predicting drug response is of a great interest to improve precision medicine. Here we conduct a comprehensive review of the main pharmacogenomics biomarkers and genomic alterations affecting enzyme activity, transporter capacity, channels and receptors, and therefore the new advances in CRC precision medicine to select the best therapeutic strategy in populations worldwide, with a focus on Latin America.

## INTRODUCTION

Colorectal cancer (CRC) is one of the leading causes of cancer death worldwide^1^. In the last decade, numerous exciting advances have been made to treat patients even with metastatic CRC^2^. However, patient-tailored therapies are still needed to overcome this disease. The advance of precision medicine requires the accurate identification of mutations driving each patient’s tumor^3^. In this regard, genetic mutations may have a great impact on disease prognosis and therapy response. Germline mutations are heritable alterations found in individuals while somatic mutations appear after an oncogenic insult within the tumoral tissue^4^. As part of CRC precision medicine, pharmacogenomics allows tailoring drug therapy based on these mutations^5^. Thus, personalized therapy not only maximizes the drug therapeutic effects but also reduces the possibility of experiencing adverse drug reactions^6^. In this review, we focus primarily on the current status of pharmacogenomics in CRC, its biomarkers and allele frequencies worldwide, with a focus on Latin American populations in order to improve precision medicine.

### Colorectal cancer oncogenomics

CRC was one of the first solid tumors to be molecularly characterized, in whose pathogenesis several signaling pathways intervene^7^. Vogelstein *et al*., described the model of progressive step-wise accumulation of epigenetic events of CRC^8–11^. This model provides information about the role of driver mutations whose objective is to give a selective advantage for tumor progression^12^. In addition, the accumulation of pathogenic mutations in the transforming growth factor-β (TGFβ), WNT-β-catenin, PI3K, EGFR and β β downstream MAPK pathways induces CRC^11, 13–15^.

On the other hand, the development of CRC also occurs when chromosomal instability (CIN) occurs, progress due to defects in telomere stability, chromosomal segregation and mutations in TP53 gene^16^. The 15% of early-stage colorectal tumors present mismatch repair-deficient (MMRd) system, triggering hypermutation and microsatellite instability (MSI)^14^. According to Dienstmann *et al*., the epigenetic profile of tumors with CIN present mutations in APC, KRAS, TP53, SMAD4 and PIK3CA, promoting the formation of the non-hypermutated consensus molecular subtypes (CMSs): CMS2, CMS3 and CMS4^1^. Whereas tumors with MSI harbor mutations in the MSH6, RNF43, ATM, TGFBR2, BRAF and PTEN genes of the hypermutated molecular subtype CMS1^1^.

### A consensus of molecular subtypes

Gene expression-based subtyping is widely accepted as a relevant source of disease stratification^17^. Nevertheless, the translational utility is hampered by divergent results that are probably related to differences in algorithms applied to sample preparation methods, gene expression platforms and racial/ethnic disparities^18, 19^. Inspection of the published gene expression-based CRC classification revealed an absence of a clear methodological ‘gold standard’^8, 9, 16, 20–23^. To facilitate clinical translation, the CRC Subtyping Consortium (CRCSC) was formed to assess the core subtype patterns among existing gene expression-based CRC subtyping algorithms^18, 24^.

In spite of heterogeneities, subtype concordance analysis readily yielded four CMSs^18^, being CMS1 the immune subtype, CMS2 the canonical subtype, CMS3 the metabolic subtype and CMS4 the mesenchymal subtype (Figure 1)^18, 25^. Upon evaluation of these classification system, Calon *et al*., discovered that their prediction power arises from genes expressed by stromal cells that associate robustly with disease relapse^26^. Mesenchymal stromal cells (MSC) may represent a pivotal part of stroma in CRC, but little is known about the specific interaction of MSC in CRC^27^.

**Figure 1.**
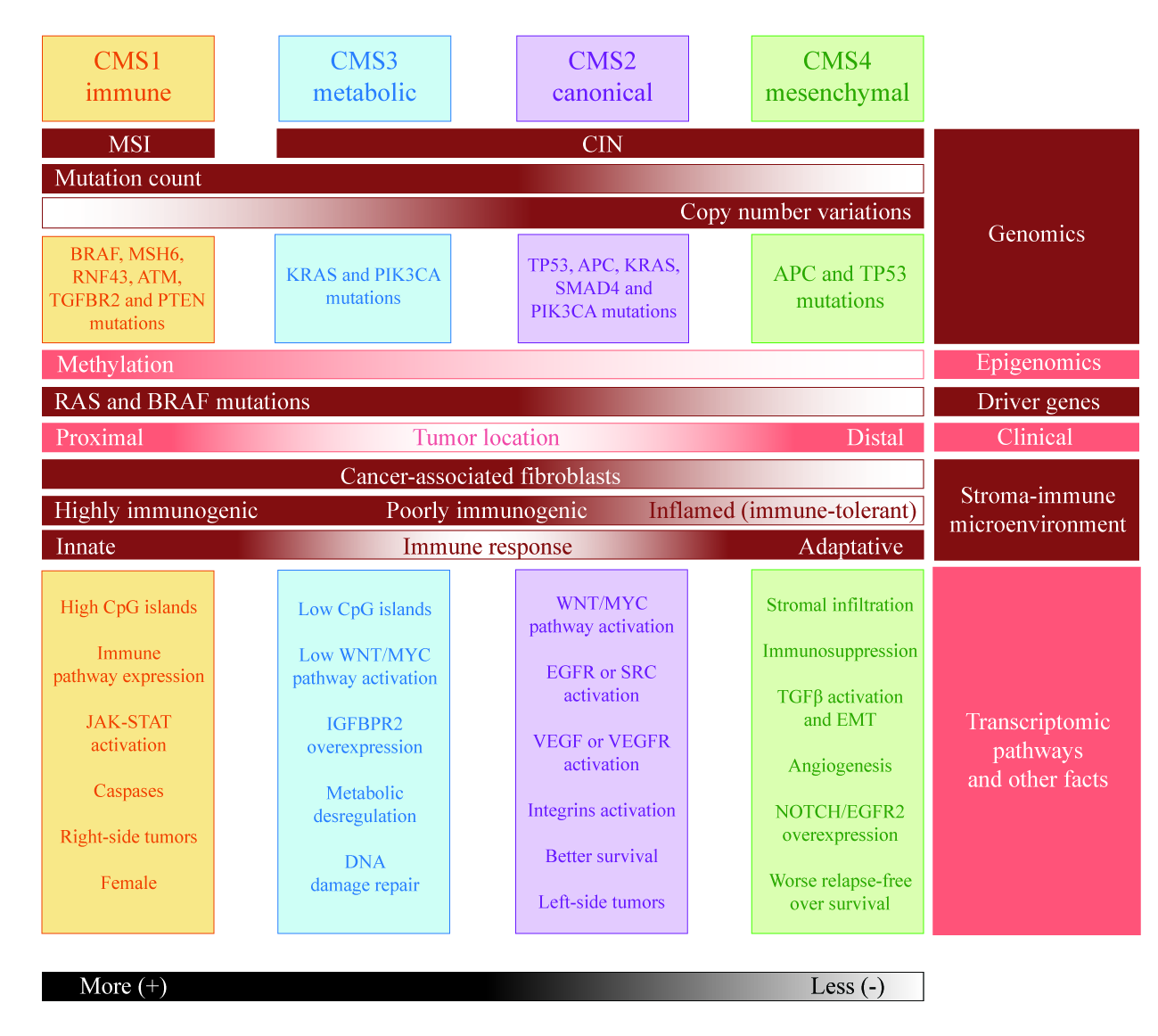
Integrating multi-omic features in CRC subtypes. Microsatellite instability (MSI) is linked to hypermutation, hypermethylation, highly immunogenic response, and locations in the proximal colon (consensus molecular subtype 1 (CMS1)). Tumors with chromosomal instability (CIN) are linked to copy number variations, poorly immunogenic or inflamed, non-hypermutated subtypes, stromal infiltration, and locations in left colon or rectum (CMS2, CMS3, and CMS4).

Recognizing that transcriptomics represents the level of high-throughput molecular data that is most intimately linked to tumor phenotype and clinical behavior, it is important to characterize the CRC genomics alterations. Tumor genomes contain thousands of mutations. However, only a few of them drive tumorigenesis by affecting driver genes, which upon alteration, confers selective growth advantage to tumor cells^28^. Since the identification of the first somatic mutation in human bladder carcinoma cell line (HRAS G12V)^29, 30^, the Pan-Cancer Atlas from The Cancer Genome Atlas (TCGA) have undertaken omics analyses identifying 20 CRC driver genes (ACVR2A, AMER1, APC, ARID1A, BRAF, CTNNB1, FBXW7, GNAS, KRAS, NRAS, PCBP1, PIK3CA, PTEN, SMAD4, SMAD2, SOX9, TCF7L2, TGIF1, TP53 and ZFP36L2) that are included in The Catalogue of Somatic Mutations in Cancer (COSMIC) Cancer Gene Census (CGC) and the Cancer Genome Interpreter (CGI)^31–35^. CGI identifies 71 biomarkers among biallelic markers, copy number alterations (CNAs), somatic mutations, fusion genes and amplifications^34^. Likewise, CGI annotates CRC tumor variants that constitute state-of-art biomarkers of drug response as shown in Supplementary Table 1.

### Drugs, biomarkers and allele frequencies

According to the National Comprehensive Cancer Network (NCCN) guidelines v1.2018 and the European Society for Medical Oncology (ESMO) guidelines^36–38^, the two main drug categories in CRC treatment are cytotoxic and biological therapies. Cytotoxic agents are platinum derivatives (oxaliplatin), antimetabolites (5-fluorouracil and capecitabine) and antitopoisomerases (irinotecan). Biological therapy includes drugs against the epidermal growth factor receptor (EGFR) (cetuximab, panitumumab) and antiangiogenics (bevacizumab, ziv-aflibercept, ramucirumab). In addition, the recommendation includes PD-1 and PD-L1 inhibitors (nivolumumab, pembrolizumab) as new immunological molecules for MSI or MMRd (Table 1 and Figure 2).

**Figure 2.**
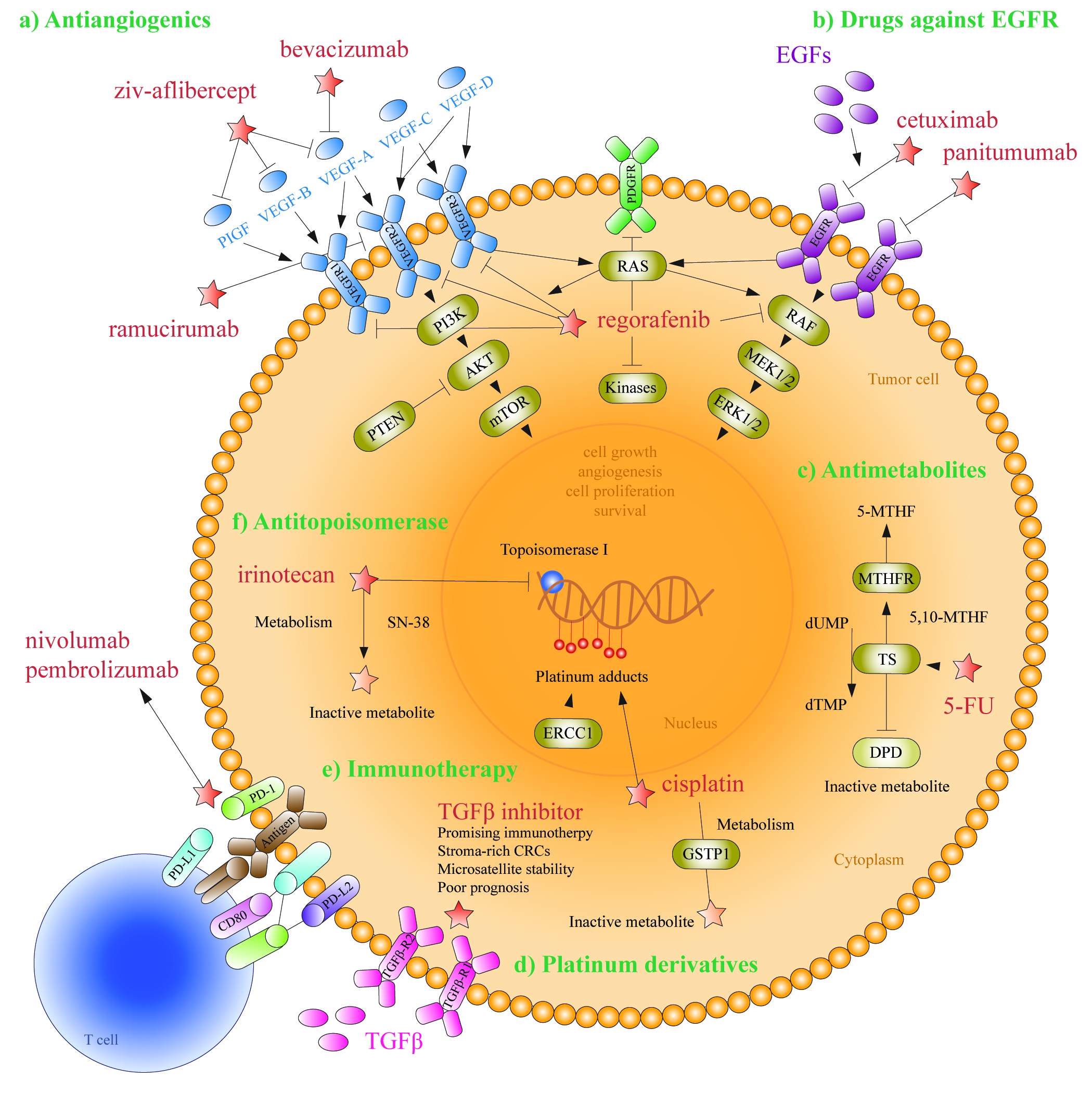
Overview of different drugs used in the CRC treatments: a) antiangiogenics, b) drugs against EGFR, c) antimetabolites, d) platinum derivatives, e) immune checkpoint inhibitors (anti PD-1, anti PD-L1 and TGF β inhibitor), and f) antitopoisomerase.

**Table 1.** Category of drugs applied in CRC treatments

| Category | Agents | Drugs |
| --- | --- | --- |
| Cytotoxic agents | Platinum derivatives | Oxaliplatin |
|  | Antimetabolites | 5-FU/leucovorin, capecitabine |
|  | Antitopoisomerase | Irinotecan |
|  | Nucleoside analogue<sup>a</sup> | Trifluridine, tripiracil |
| Monoclonal antibodies | Antiangiogenic<sup>b</sup> | Bevacizumab, ziv-aflibercept, ramucirumab |
|  | Anti EGFR<sup>b</sup> | Cetuximab, panitumumab |
|  | Anti BRAF V600E<sup>b</sup> | Vemurafenib |
|  | Anti PD-L1, PD-1<sup>b</sup> | Nivolumab, pembrolizumab |
| Tyrosine kinase inhibitors<sup>a</sup> | Poli anti kinase inhibitors | Regorafenib, sorafenib |

5-FU, 5-fluorouracil; EGFR, epidermal growth factor receptor; BRAF, B-Raf protooncogene; PD-1, programmed cell death protein 1; PD-L1, programmed cell death ligand 1. <sup>a</sup>For patients who have progressed through all available regimens. <sup>b</sup>For advanced or metastatic disease only.

#### Platinum derivatives

These compounds form covalent bonds with guanine and adenine in the DNA. The most important drugs in this group are cisplatin, carboplatin and oxaliplatin (Figure 2). Drugs containing platinum salts exert their cytotoxic effect by means of DNA adduct formation, leading to inhibition of DNA replication and apoptosis^39^. The major path of adduct elimination is the nucleotide excision repair (NER). During NER, damaged DNA and unwound DNA helices are identified by the action of several factors, including xeroderma pigmentosum proteins (XPD, XPC and XPA). Cleavages of the damaged DNA strand are performed by nucleases XPG (3’) and ERCC1 (5’), and adducts are removed^40^.

The glutathione S-transferases (GSTs) are involved in the inactivation of platinum compounds, thus preventing cellular DNA damage and increasing the treatment efficacy^41^. Single nucleotide polymorphisms (SNPs) in GSTP1, GSTT1 and GSTM1 can alter GST activity^42^. The decrease in enzyme activity has been linked to reduced detoxification capacity, leading to increased efficacy of platinum compounds. GSTP1 (Ile105Val) has been associated with reduced enzyme activity43, a deletion in GSTT1 leads to the absence of enzyme activity and a deletion in GSTM1 is linked with decreased survival rate^44, 45^.

The excision repair cross-complementation (ERCC) is involved in nucleotide repair system^46^. Polymorphisms in genes encoding these repair proteins may contribute to inter-individual differences to platinum toxicity. The association between toxicity and ERCC1 rs11615 has been studied in CRC^47^. The mutant T allele has been related to grade 1 neuropathy in oxaliplatin-treated patients, even though no association with a higher degree of neuropathy was observed. In addition, ERCC2 (XPD) is involved in the oxaliplatin pathway. rs13181 has been related with treatment effectiveness^48^. Meanwhile, ERCC4 rs1799801, ERCC5 rs2016073 and rs751402 are associated with platinum response^49^.

The X-ray repair cross-complementing protein (XRCC1) and its variant rs25487 are involved in the repair of broken DNA strands which can be induced by platinum compounds; such repair is carried out by an excision repair system^50^. It has been suggested that a deterioration in the efficiency of DNA repair caused by Gln’s allele leads to greater efficacy of oxaliplatin^50^. On the other hand, the presence of XRCC3 rs1799794 has been associated with an increased risk of neutropenia^51^. Biomarkers focused on oxaliplatin are listed in Table 2^42, 48–50, 52–55^.

**Table 2.** Biomarkers focused on oxaliplatin

| Gene | Polymorphism | Clinical relevance | Function | Type of inheritance | Reference |
| --- | --- | --- | --- | --- | --- |
| GSTP1 | rs1695 (A313G) | Neurotoxicity, neutropenia | Enzyme | Germinal | <sup>50,52</sup> |
| GSTM1 | Del | Poor survival, neutropenia | Enzyme | Germinal | <sup>52</sup> |
| ERCC1 | rs11615 (T354C) | Neuropathy / Survival | Repair protein | Germinal | <sup>50</sup> |
| ERCC2 | rs13181 (A2251C/T) | Effectiveness / Survival | Repair protein | Germinal | <sup>42,48</sup> |
| ERCC4 | rs1799801 (T2505C) | Response | Repair protein | Germinal | <sup>53</sup> |
| ERCC5 | rs2016073 (A-763G);<br>rs751402 (A+25G) | Response | Repair protein | Germinal | <sup>49</sup> |
| XRCC1 | rs25487 (G1196A) | Response | Repair protein | Somatic | <sup>50,54</sup> |
| XRCC3 | rs1799794 (A316G) | Neutropenia | Repair protein | Germinal | <sup>55</sup> |

GSTP1, glutathione S-transferase pi 1; GSTM1, glutathione S-transferase mu 1; ERCC1, excision repair 1; ERCC2, excision repair 2; ERCC4, excision repair 4; ERCC5, excision repair 5; XRCC1, X-ray repair cross complementing 1; XRCC3, X-ray repair cross complementing 3; del, deletion; G, guanine; A, adenine; C, cytosine; T, thymine.

Pharmacogenomics identifies mutations that may predict an efficient therapeutic response; however, genetic variations significantly change among race/ethnic populations worldwide^56^. The allele frequencies of platinum derivative variants rs1695, rs11615, rs13181, rs1799801, rs2016073, rs2234671, rs25487 and rs1799794, according to the 1000 Genomes Project (phase 3), are shown in Table 3^57^.

**Table 3.** Allele frequencies for clinically relevant genetic variants GSTP1 rs1695, ERCC1 rs11615, ERCC2 rs13181, ERCC4 rs1799801, ERCC5 rs2016073, CXCR1 rs2234671, XRCC1 rs25487 and XRCC3 rs1799794 in populations worldwide

| Gene | Polymorphism | Human Populations |  |  |
| --- | --- | --- | --- | --- |
|  |  | Latin American | Caucasian | Asian |
| GSTP1 | rs1695<br>(A313G) | Colombia: 0.36 (G)<sup>a</sup>;<br>Mexico: 0.56 (G);<br>Peru: 0.67 (G);<br>Puerto Rico: 0.37 (G) | Spain: 0.36 (G);<br>British: 0.32 (G);<br>Finland: 0.28 (G);<br>Italy: 0.29 (G) | Han Chinese: 0.18 (G);<br>Bangladesh: 0.22 (G);<br>Japan: 0.10 (G);<br>Vietnam: 0.20 (G) |
| ERCC1 | rs11615<br>(T354C) | Colombia: 0.52 (G);<br>Mexico: 0.74 (G);<br>Peru: 0.75 (G);<br>Puerto Rico: 0.50 (G) | Spain: 0.37 (G);<br>British: 0.32 (G);<br>Finland: 0.37 (G);<br>Italy: 0.46 (G) | Han Chinese: 0.75 (G);<br>Bangladesh: 0.65 (G);<br>Japan: 0.71 (G);<br>Vietnam: 0.73 (G) |
| ERCC2 | rs13181<br>(A2251C/T) | Colombia: 0.24 (G);<br>Mexico: 0.19 (G);<br>Peru: 0.17 (G);<br>Puerto Rico: 0.24 (G) | Spain: 0.31 (G);<br>British: 0.30 (G);<br>Finland: 0.40 (G);<br>Italy: 0.45 (G) | Han Chinese: 0.11 (G);<br>Bangladesh: 0.35 (G);<br>Japan: 0.07 (G);<br>Vietnam: 0.08 (G) |
| ERCC4 | rs1799801<br>(T2505C) | Colombia: 0.22 (G);<br>Mexico: 0.19 (G);<br>Peru: 0.24 (G);<br>Puerto Rico: 0.19 (G) | Spain: 0.38 (G);<br>British: 0.28 (G);<br>Finland: 0.25 (G);<br>Italy: 0.29 (G) | Han Chinese: 0.21 (G);<br>Bangladesh: 0.24 (G);<br>Japan: 0.33 (G);<br>Vietnam: 0.37 (G) |
| ERCC5 | rs2016073<br>(A-763G) | Colombia: 0.73 (A);<br>Mexico: 0.61 (A);<br>Peru: 0.48 (A);<br>Puerto Rico: 0.78 (A) | Spain: 0.87 (A);<br>British: 0.80 (A);<br>Finland: 0.82 (A);<br>Italy: 0.77 (A) | Han Chinese: 0.65 (A);<br>Bangladesh: 0.66 (A);<br>Japan: 0.73 (A);<br>Vietnam: 0.63 (A) |
| CXCR1 | rs2234671<br>(G2607C) | Colombia: 0.08 (G);<br>Mexico: 0.14 (G);<br>Peru: 0.31 (G);<br>Puerto Rico: 0.08 (G) | Spain: 0.01 (G);<br>British: 0.08 (G);<br>Finland: 0.04 (G);<br>Italy: 0.03 (G) | Han Chinese: 0.10 (G);<br>Bangladesh: 0.18 (G);<br>Japan: 0.08 (G);<br>Vietnam: 0.06 (G) |
| XRCC1 | rs25487<br>(G1196A) | Colombia: 0.63 (C);<br>Mexico: 0.73 (C);<br>Peru: 0.69 (C);<br>Puerto Rico: 0.71 (C) | Spain: 0.58 (C);<br>British: 0.66 (C);<br>Finland: 0.67 (C);<br>Italy: 0.63 (C) | Han Chinese: 0.75 (C);<br>Bangladesh: 0.66 (C);<br>Japan: 0.72 (C);<br>Vietnam: 0.77 (C) |
| XRCC3 | rs1799794<br>(A316G) | Colombia: 0.19 (C);<br>Mexico: 0.16 (C);<br>Peru: 0.25 (C);<br>Puerto Rico: 0.16 (C) | Spain: 0.28 (C);<br>British: 0.19 (C);<br>Finland: 0.23 (C);<br>Italy: 0.19 (C) | Han Chinese: 0.52 (C);<br>Bangladesh: 0.42 (C);<br>Japan: 0.42 (C);<br>Vietnam: 0.39 (C) |

GSTP1, glutathione S-transferase pi 1; ERCC1, excision repair 1; ERCC2, excision repair 2; ERCC4, excision repair 4; ERCC5, excision repair 5; XRCC1, X-ray repair cross complementing 1; XRCC3, X-ray repair cross complementing 3; CXCR1, C-X-C motif chemokine receptor 1; G, guanine; A, adenine; C, cytosine; T, thymine. <sup>a</sup> Frequency of minor allele.

#### Antimetabolites

These drugs inhibit enzymes related to purine and pyrimidine synthesis, resulting in cell depletion and alteration of nucleic acid synthesis. Among these, there are pyrimidine analogs such as 5-fluorouracil (5-FU) and oral pro-drugs such as gemcitabine, capecitabine and tegafur (Figure 2)^58^.

Fluoropyrimidines (5-FU, capecitabine and tegafur) are antimetabolite drugs used in CRC treatment. 5-FU is a fluoropyrimidine derivative with two major mechanisms of action that explain its cytotoxic effect^59^. The main active metabolite of 5-FU (5-FdUMP) prevents DNA synthesis by forming a complex with thymidylate synthase (TS) stabilized by 5,10-methylenetetrahydrofolate (5,10-MTHF), thus inhibiting the conversion of monophosphate 2’-deoxyuridine-5’ (dUMP) to deoxythymidine-2’-5’-monophosphate (dTMP), an essential precursor for DNA synthesis. In addition, the incorporation of 5-FU to nucleotides in DNA and RNA strands leads to an alteration in the processing of nucleic acids^59^. Gemcitabine is a structural analog of deoxycytidine, which is metabolized by nucleoside kinase to nucleoside diphosphate and triphosphate^60^.

TYMS protein is a homodimeric methyltransferase enzyme that catalyzes the synthesis reaction of thymidylate. This reaction is a critical step in the formation of deoxythymidine 5’-triphosphate (dTTP), an indispensable metabolite in DNA synthesis. TYMS contains a tandem of a polymorphic 28-base-pair sequence repeated in the promoter (TSER) 5’ untranslated region (5’-UTR)^42, 60^.

Inactivation of 5-FU depends on dihydropyrimidine dehydrogenase (DPYD) activity^61^. The deficient activity of DPYD leads to prolonged 5-FU plasma half-life, causing a severe hematological toxicity^62^. DPYD deficiency is present in ∼3% of all cancer patients, but it represents approximately 50% of patients manifesting severe toxicity. So far, ∼30 polymorphisms in DYPD have been identified; however, a mutation of G>A at the splicing site in exon 14 (IVS14+1G>A) leads to the formation of a truncated protein without residual activity^63^. The incidence of this allele is rare, with a heterozygote population frequency of 0.9-1.8%. Nevertheless, it is estimated to be responsible for approximately 25% of all cases of 5-FU unexpected toxicity^59^.

Methylenetetrahydrofolate reductase (MTHFR) catalyzes the conversion of 5,10-MTHF to 5-MTHF. The most common polymorphisms of MTHFR are C677T and A1298C. These polymorphisms lead to decreased enzyme activity, which induces a more effective stabilization of the FdUMP-TS ternary complex, potentiating 5-FU toxicity^64–66^.

ATP-Binding Cassette Sub-Family B1 (ABCB1) is a member of the ABC transporter superfamily and its protein is known as P-glycoprotein^67^. ABCB1 overexpression in tumors has been associated with resistance to chemotherapeutic drugs. ABCB1 is a highly polymorphic gene that significantly differs among ethnic groups. Some of the most studied SNPs are rs1128503, rs2032592 and rs1045642^68^.

Cytidine deaminase (CDA) is involved in capecitabine metabolism in the liver to form 5-fluorodeoxyuridine, which, in turn, becomes 5-FU by the action of thymidine phosphorylase (TP)^60^. The decreased activity of CDA leads to the accumulation of potentially toxic metabolites. The variation in expression of CDA has been linked to polymorphisms in the promoter region of CDA and affects the metabolism of gemcitabine and capecitabine^67^. rs602950 and rs532545 have been associated with increased expression of CDA *in vitro* in capecitabine-treated patients^67^.

TP is involved in 5-FU metabolism, where 5-FU is converted to 5-fluoro-2’-deoxyuridine (FUDR-5)^69^. rs11479 generates an amino acid change of serine to leucine that leads to a lower treatment response. Enolase superfamily member 1 (ENOSF1) gene encodes an antisense RNA against TYMS. ENOSF1 regulates mRNA and protein expression of TYMS. Hence, TYMS variants with lower or higher activity affect its function^63, 64^. Lastly, biomarkers focused on capecitabine, 5-FU and gemcitabine drugs are listed in Table 4 ^63, 64, 67–73^.

**Table 4.** Biomarkers focused on capecitabine, 5-FU and gemcitabine drugs

| Gene | Polymorphism | Clinical relevance | Function | Type of inheritance | Reference |
| --- | --- | --- | --- | --- | --- |
| TYMS | rs45445694 (*2R/*3R) | Neutropenia | Enzyme | Germinal | <sup>63,70</sup> |
| DPYD | rs3918290 (G1905+1A);<br>rs67376798 (A2846T);<br>rs1801158 (G1601A);<br>rs55886062 (T1679G);<br>rs1801159 (A1627G);<br>rs12132152 (G97057448A);<br>rs12022243 (C97397224T) | Toxicity<br>Toxicity<br>Toxicity<br>Toxicity<br>Neutropenia<br>Diarrhea<br>Diarrhea | Enzyme | Germinal | <sup>63,64,71</sup> |
| MTHFR | rs1801131 (A1298T);<br>rs1801133 (C677T) | Toxicity<br>Toxicity | Enzyme | Germinal | <sup>70,72</sup> |
| ABCB1 | rs1128503 (C1236T);<br>rs1045642 (C3435T) | Neutropenia<br>Diarrhea | Transporter | Germinal | <sup>67,68</sup> |
| CDA | rs2072671 (A79C);<br>rs602950 (A-92G);<br>rs532545 (C-451T) | Toxicity<br>Diarrhea<br>Diarrhea | Enzyme | Germinal | <sup>67</sup> |
| CCND1 | rs9344 (G870A) | Prognosis | Cyclin | Germinal | <sup>73</sup> |
| TP | rs11479 (C1412T) | Response | Enzyme | Germinal | <sup>69</sup> |
| ENOSF1 | rs2612091 (805-227G>A) | Diarrhea | Enzyme | Germinal | <sup>64,67</sup> |

ABCB1, ATP binding cassette subfamily B member 1; CCND1, cyclin D1; CDA, cytidine deaminase; DPYD, dihydropyrimidine dehydrogenase; ENOSF1, enolase superfamily member 1; MTHFR, methylenetetrahydrofolate reductase; TP, thymidine phosphorylase; TYMS, thymidylate synthetase; G, guanine; A, adenine; C, cytosine; T, thymine.

The allele frequencies of rs3918290, rs1801133, rs1128503, rs2072671, rs9344, rs9344 and rs2612091 polymorphisms in populations worldwide are shown in Table 5^57^.

**Table 5.** Allele frequencies for clinically relevant genetic variants DPYD rs3918290, MTHFR rs1801133, ABCB1 rs1128503, CDA rs2072671, CCND1 rs9344, TP rs9344 and ENOSF1 rs2612091 in populations worldwide

| Gene | Polymorphism | Human Populations |  |  |
| --- | --- | --- | --- | --- |
|  |  | Latin American | Caucasian | Asian |
| DPYD | rs3918290<br>(1905+1G>A) | Colombia: 0.00 (T)<sup>a</sup>;<br>Mexico: 0.00 (T);<br>Peru: 0.01 (T);<br>Puerto Rico: 0.00 (T) | Spain: 0.00 (T);<br>British: 0.00 (T);<br>Finland: 0.02 (T);<br>Italy: 0.00 (T) | Han Chinese: 0.00 (T);<br>Bangladesh: 0.00 (T);<br>Japan: 0.00 (T);<br>Vietnam: 0.00 (T) |
| MTHFR | rs1801133<br>(C677T) | Colombia: 0.54 (A);<br>Mexico: 0.47 (A);<br>Peru: 0.44 (A);<br>Puerto Rico: 0.45 (A) | Spain: 0.44 (A);<br>British: 0.32 (A);<br>Finland: 0.27 (A);<br>Italy: 0.47 (A) | Han Chinese: 0.47 (A);<br>Bangladesh: 0.12 (A);<br>Japan: 0.38 (A);<br>Vietnam: 0.19 (A) |
| ABCB1 | rs1128503<br>(C1236T) | Colombia: 0.57 (G);<br>Mexico: 0.53 (G);<br>Peru: 0.67 (G);<br>Puerto Rico: 0.60 (G) | Spain: 0.62 (G);<br>British: 0.58 (G);<br>Finland: 0.57 (G);<br>Italy: 0.58 (G) | Han Chinese: 0.30 (G);<br>Bangladesh: 0.37 (G);<br>Japan: 0.40 (G);<br>Vietnam: 0.42 (G) |
| CDA | rs2072671<br>(A79C) | Colombia: 0.27 (C);<br>Mexico: 0.32 (C);<br>Peru: 0.36 (C);<br>Puerto Rico: 0.28 (C) | Spain: 0.34 (C);<br>British: 0.33 (C);<br>Finland: 0.19 (C);<br>Italy: 0.37 (C) | Han Chinese: 0.12 (C);<br>Bangladesh: 0.17 (C);<br>Japan: 0.21 (C);<br>Vietnam: 0.10 (C) |
| CCND1 | rs9344<br>(G870A) | Colombia: 0.33 (A);<br>Mexico: 0.33 (A);<br>Peru: 0.30 (A);<br>Puerto Rico: 0.42 (A) | Spain: 0.55 (A);<br>British: 0.47 (A);<br>Finland: 0.42 (A);<br>Italy: 0.51 (A) | Han Chinese: 0.56 (A);<br>Bangladesh: 0.58 (A);<br>Japan: 0.47 (A);<br>Vietnam: 0.63 (A) |
| TP | rs11479<br>(C1412T) | Colombia: 0.11 (A);<br>Mexico: 0.25 (A);<br>Peru: 0.24 (A);<br>Puerto Rico: 0.12 (A) | Spain: 0.06 (A);<br>British: 0.05 (A);<br>Finland: 0.08 (A);<br>Italy: 0.07 (A) | Han Chinese: 0.24 (A);<br>Bangladesh: 0.16 (A);<br>Japan: 0.25 (A);<br>Vietnam: 0.32 (A) |
| ENOSF1 | rs2612091<br>(805-227G>A) | Colombia: 0.61 (T);<br>Mexico: 0.59 (T);<br>Peru: 0.69 (T);<br>Puerto Rico: 0.61 (T) | Spain: 0.54 (T);<br>British: 0.53 (T);<br>Finland: 0.57 (T);<br>Italy: 0.56 (T) | Han Chinese: 0.70 (T);<br>Bangladesh: 0.55 (T);<br>Japan: 0.69 (T);<br>Vietnam: 0.70 (T) |

ABCB1, ATP binding cassette subfamily B member 1; CCND1, cyclin D1; CDA, cytidine deaminase; DPYD, dihydropyrimidine dehydrogenase; ENOSF1, enolase superfamily member 1; MTHFR, methylenetetrahydrofolate reductase; TP, thymidine phosphorylase; G, guanine; A, adenine; C, cytosine; T, thymine. <sup>a</sup> Frequency of minor allele.

#### Agents interacting with topoisomerases

Topoisomerases play a key role in the cell replication, transcription, and DNA repair. It modifies the tertiary DNA structure without altering the nucleotide sequence. In humans, three types of topoisomerases (I, II and III) have been identified. Within this group, camptothecin derivatives are included such as irinotecan^74^ (Figure 2).

Irinotecan is a potent inhibitor of topoisomerase I^75^. It promotes an oxidation bioreaction mediated by CYP3A to form APC, a cytotoxic substance. Alternatively, irinotecan is converted by hepatic carboxylesterase to SN-38. This compound is conjugated further by several UDP glucuronyl to reach the inactive metabolite SN-38G^76^. To enable excretion, SN-38 and irinotecan are actively transported out of the cell by ATP-dependent efflux pump (ABCB1). After biliary excretion, SN-38G can become active SN-38 by bacterial beta-glucuronidase, which can lead to gastrointestinal toxicity.

It has been shown that reduced glucuronidation of SN-38 significantly increases irinotecan gastrointestinal toxicity^77^. The main UDP-glucuronosyltransferase (UGT) involved in conjugating SN-38 is UGT1A1. At least 25 UGT1A1 polymorphisms have been described, of which the most common in the promoter region consists of seven TA-repetitions (−53 [TA] 6>7, UGT1A1*28) instead of six^78, 79^. The highest number of TA repeats is associated with a reduction of UGT1A1 expression, leading to reduced glucuronidation. UGT1A1*28 has proven to be a significant predictor of severe toxicity following administration of irinotecan^80, 81^.

ABC transporters, including ABCC1, ABCC2, ABCB1 and ABCG2, regulate output of hepatic and biliary CPT-11 metabolites^55, 82^. SNPs in ABCB1 and ABCC2 have been recently associated with modulation of CPT-11 and SN-38 exposure^75^. In addition, other SNPs in ABCC5 and ABCG2 genes have been correlated with both hematological and non-hematological toxicities^83^.

The solute carrier organic anion transporter family member 1B1 (SLCO1B1) is an important transporter expressed in the basolateral membrane of hepatocytes which mediates the availability of active irinotecan metabolite^84^. rs4149056 has been associated with an increased SN-38 concentration in patients with metastatic CRC (mCRC). Meanwhile, other polymorphisms are linked to faster response rate and higher PFS^76^. Finally, biomarkers focused on irinotecan drug are shown in Table 6^45, 52, 68, 78, 79, 81, 83–89^.

**Table 6.** Biomarkers focused on irinotecan drug

| Gene | Polymorphism | Clinical relevance | Function | Type of inheritance | Reference |
| --- | --- | --- | --- | --- | --- |
| CYP3A5 | rs776746 (*3C) | Response | Enzyme | Germinal | 45 |
| UGT1A1 | rs8175347 (*28);<br>rs4148323 (*6) | Neutropenia and Diarrhea<br>Neutropenia | Enzyme | Germinal | 78,79,81 |
| UGT1A7 | rs17868324 (*3) | Neutropenia | Enzyme | Germinal | 79,81 |
| UGT1A9 | rs3832043 (*22) | Neutropenia | Enzyme | Germinal | 79,81,88 |
| ABCB1 | rs1128503 (C1236T);<br>rs1045642 (C3435T);<br>rs2032582 (G2677T/A) | Asthenia<br>Diarrhea<br>Global Survival | Transporter | Germinal | 67,68 |
| ABCC1 | rs2074087 (C2461-30G) | Neutropenia | Transporter | Germinal | 52,55 |
| ABCC2 | rs3740066 (T3972C) | Neutropenia | Transporter | Germinal | 86,87,89 |
| ABCG2 | rs262604 (-20+805 A>G);<br>rs2231142 (C421A);<br>rs7699188 (C61414T) | Myelosuppression<br>Neutropenia<br>Toxicity | Transporter | Germinal | 83 |
| SLC19A1 | rs1051266 (A80G) | PFS | Transporter | Germinal | 84 |
| SLCO1B1 | rs2306283 (A388G) | PFS | Transporter | Germinal | 84 |

ABCB1, ATP binding cassette subfamily B member 1; ABCC1, ATP binding cassette subfamily C member 1; ABCC2, ATP binding cassette subfamily C member 2; ABCG2, ATP binding cassette subfamily G member 2; PFS, progression-free survival; CYP3A5, cytochrome P450 family 3 subfamily A member 5; UGT1A1, UDP glucuronosyltransferase family 1 member A1; UGT1A7, UDP glucuronosyltransferase family 1 member A7; UGT1A9, UDP glucuronosyltransferase family 1 member A9; SLC19A1, solute carrier family 19 member 1; SLCO1B1, solute carrier organic anion transporter family member 1B1; G, guanine; A, adenine; C, cytosine; T, thymine; \*, repetitions

The allele frequencies of genes that interact with topoisomerases rs2244613, rs1045642, rs2074087, rs262604, rs1051266 and rs2306283 in populations worldwide are shown in Table 7^57^.

**Table 7.**
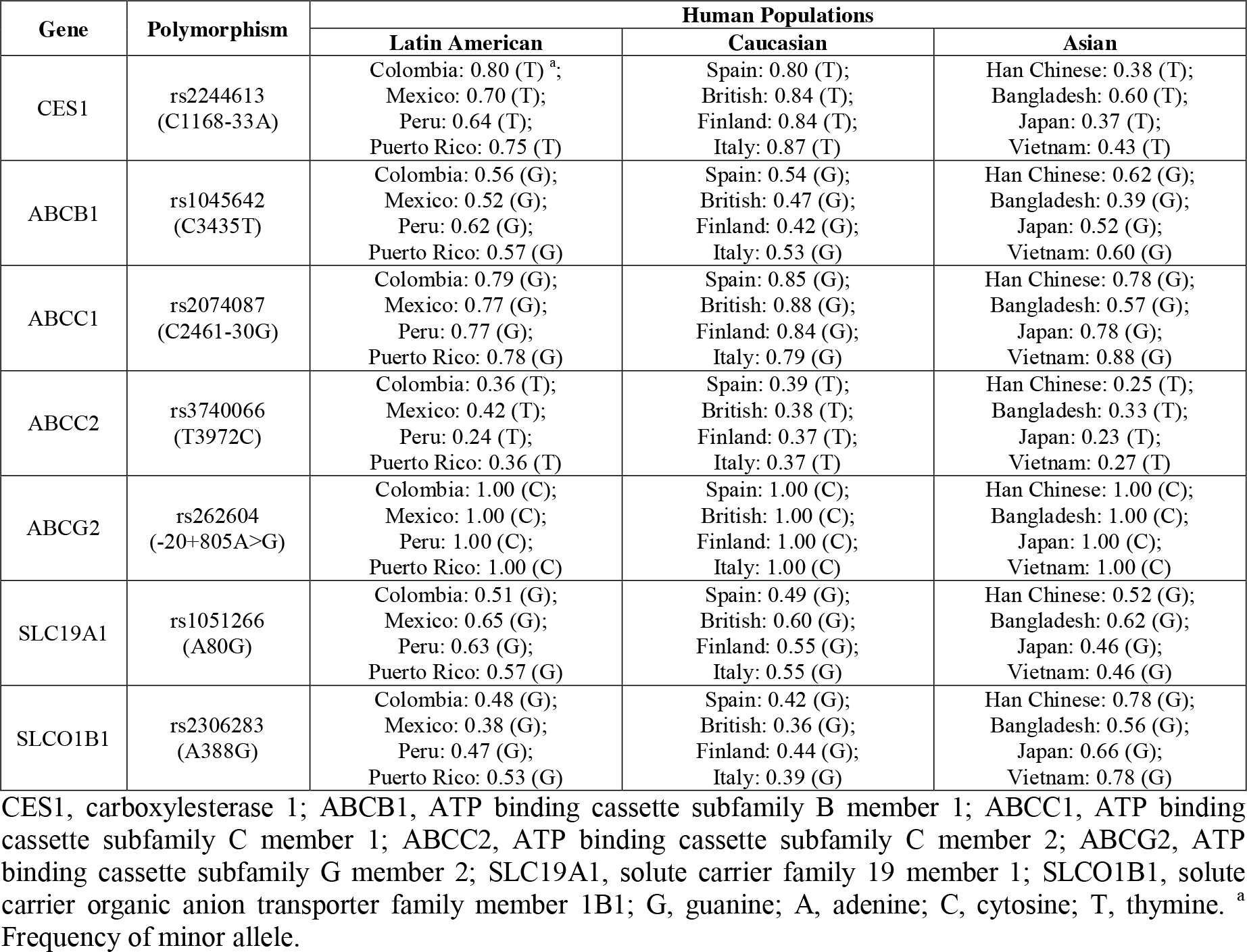
Allele frequencies for clinically relevant germline polymorphisms CES1 rs2244613, ABCB1 rs1045642, ABCC1 rs2074087, ABCG2 rs262604, SCL19A1 rs1051266 and SLCO1B1 rs2306283 in populations worldwide

| Gene | Polymorphism | Human Populations |  |  |
| --- | --- | --- | --- | --- |
|  |  | Latin American | Caucasian | Asian |
| CES1 | rs2244613<br>(C1168-33A) | Colombia: 0.80 (T)<sup>a</sup>;<br>Mexico: 0.70 (T);<br>Peru: 0.64 (T);<br>Puerto Rico: 0.75 (T) | Spain: 0.80 (T);<br>British: 0.84 (T);<br>Finland: 0.84 (T);<br>Italy: 0.87 (T) | Han Chinese: 0.38 (T);<br>Bangladesh: 0.60 (T);<br>Japan: 0.37 (T);<br>Vietnam: 0.43 (T) |
| ABCB1 | rs1045642<br>(C3435T) | Colombia: 0.56 (G);<br>Mexico: 0.52 (G);<br>Peru: 0.62 (G);<br>Puerto Rico: 0.57 (G) | Spain: 0.54 (G);<br>British: 0.47 (G);<br>Finland: 0.42 (G);<br>Italy: 0.53 (G) | Han Chinese: 0.62 (G);<br>Bangladesh: 0.39 (G);<br>Japan: 0.52 (G);<br>Vietnam: 0.60 (G) |
| ABCC1 | rs2074087<br>(C2461-30G) | Colombia: 0.79 (G);<br>Mexico: 0.77 (G);<br>Peru: 0.77 (G);<br>Puerto Rico: 0.78 (G) | Spain: 0.85 (G);<br>British: 0.88 (G);<br>Finland: 0.84 (G);<br>Italy: 0.79 (G) | Han Chinese: 0.78 (G);<br>Bangladesh: 0.57 (G);<br>Japan: 0.78 (G);<br>Vietnam: 0.88 (G) |
| ABCC2 | rs3740066<br>(T3972C) | Colombia: 0.36 (T);<br>Mexico: 0.42 (T);<br>Peru: 0.24 (T);<br>Puerto Rico: 0.36 (T) | Spain: 0.39 (T);<br>British: 0.38 (T);<br>Finland: 0.37 (T);<br>Italy: 0.37 (T) | Han Chinese: 0.25 (T);<br>Bangladesh: 0.33 (T);<br>Japan: 0.23 (T);<br>Vietnam: 0.27 (T) |
| ABCG2 | rs262604<br>(-20+805A>G) | Colombia: 1.00 (C);<br>Mexico: 1.00 (C);<br>Peru: 1.00 (C);<br>Puerto Rico: 1.00 (C) | Spain: 1.00 (C);<br>British: 1.00 (C);<br>Finland: 1.00 (C);<br>Italy: 1.00 (C) | Han Chinese: 1.00 (C);<br>Bangladesh: 1.00 (C);<br>Japan: 1.00 (C);<br>Vietnam: 1.00 (C) |
| SLC19A1 | rs1051266<br>(A80G) | Colombia: 0.51 (G);<br>Mexico: 0.65 (G);<br>Peru: 0.63 (G);<br>Puerto Rico: 0.57 (G) | Spain: 0.49 (G);<br>British: 0.60 (G);<br>Finland: 0.55 (G);<br>Italy: 0.55 (G) | Han Chinese: 0.52 (G);<br>Bangladesh: 0.62 (G);<br>Japan: 0.46 (G);<br>Vietnam: 0.46 (G) |
| SLCO1B1 | rs2306283<br>(A388G) | Colombia: 0.48 (G);<br>Mexico: 0.38 (G);<br>Peru: 0.47 (G);<br>Puerto Rico: 0.53 (G) | Spain: 0.42 (G);<br>British: 0.36 (G);<br>Finland: 0.44 (G);<br>Italy: 0.39 (G) | Han Chinese: 0.78 (G);<br>Bangladesh: 0.56 (G);<br>Japan: 0.66 (G);<br>Vietnam: 0.78 (G) |

CES1, carboxylesterase 1; ABCB1, ATP binding cassette subfamily B member 1; ABCC1, ATP binding cassette subfamily C member 1; ABCC2, ATP binding cassette subfamily C member 2; ABCG2, ATP binding cassette subfamily G member 2; SLC19A1, solute carrier family 19 member 1; SLCO1B1, solute carrier organic anion transporter family member 1B1; G, guanine; A, adenine; C, cytosine; T, thymine. <sup>a</sup> Frequency of minor allele.

#### Antiangiogenics

Vascular endothelial growth factor (VEGF) is a major regulator of angiogenesis and inhibition mediated by bevacizumab which reduces tumor volume^90, 91^. Bevacizumab is a recombinant humanized monoclonal IgG antibody directed against all isoforms of VEGF-A. Hypoxia is a potent stimulus for VEGF expression and one of the control elements in this mechanism is the hypoxia-inducible factor 1A (HIF-1a)^92^. This factor binds to a 28-bp promoter in the 5’ upstream region of VEGF, thereby/hence-stimulating transcription. In addition, other regulatory elements for VEGF expression are found in the 3’-UTR as shown in Table 8^92–95^.

**Table 8.** Biomarkers focused on antiangiogenics and bevacizumab drug

| Gene | Polymorphism | Clinical relevance | Function | Type of inheritance | Reference |
| --- | --- | --- | --- | --- | --- |
| VEGFR1 | rs9582036 (C-834A) | OS | Receptor | Germinal | <sup>94</sup> |
| VEGFR2 | rs12505758 (2266+1166 A>G) | PFS | Receptor | Germinal | <sup>92,94</sup> |
| VEGFA | rs3025039 (C*237T) | PFS | Growth factor | Germinal | <sup>94</sup> |
|  | rs13207351 (G-152A) | PFS |  | Germinal |  |
| ANXA11 | rs1049550 (C688T) | ORR | Calcium-dependent phospholipid-binding proteins | Germinal | <sup>95</sup> |
| CXCR1 | rs2234671 (G827C) | Response | Receptor | Germinal | <sup>93</sup> |
| CXCR2 | rs2230054 (C786T) | ORR | Receptor | Germinal | <sup>93</sup> |

ANXA11, annexin A11; CXCR1, C-X-C motif chemokine receptor 1; CXCR2, C-X-C motif chemokine receptor 2; VEGFA, vascular endothelial growth factor A; VEGFR1, vascular endothelial growth factor receptor 1; VEGFR2, vascular endothelial growth factor receptor 2; G, guanine; A, adenine; C, cytosine; T, thymine; OS, overall survival; PFS, patient-free survival; ORR, overall response rates.

Variations in the VEGF receptor 1 (rs9582036) and 2 (rs12505758) are associated with tyrosine kinase domain. High expression levels of these receptors contribute to a less favorable outcome when treated with bevacizumab, associated with PFS and overall survival (OS)^90, 94, 96^.

Various studies from mCRC have investigated the predictive impact of some SNPs present in VEGF-A, which are involved in bevacizumab response. Loupakis *et al*., conducted a retrospective analysis which found that rs833061 was associated with PFS and OS^94^. Meanwhile, Sibertin-Blanc *et al*., showed that T-carriers of the C237T SNP had shorter time-to-treatment failure as well as shorter PFS and OS^97^.

Annexin A11 (ANXA11) has been associated with a spectrum of regulatory functions in calcium signaling, cell division and apoptosis^98^. ANXA11 rs1049550 leads to an amino acid change (R230C) of the first conserved domain of annexin, which is responsible for Ca^+2^ dependent intracellular traffic. Response to bevacizumab revealed that patients carrying rs1049550 were more sensitive to chemotherapy than those having at least one C allele^95^.

CXC chemokine receptors (CXCR1 and CXCR2) are integral membrane proteins which specifically bind and respond to CXC chemokine family cytokines^93^. They represent a family of seven receptors linked to G-protein that plays an important role in angiogenesis. CXCR1 rs2234671 and CXCR2 rs2230054 are associated with overall response (ORR)^99^.

Finally, the Food and Drug Administration (FDA) and the European Medicines Agency (EMA) have approved new molecules that improve the therapeutic effectiveness, OS and PFS. In particular, research efforts have focused on novel agents targeting tumor angiogenic activity, cell growth and migration in mCRC. The use of molecules targeting VEGF pathways (ziv-aflibercept, regorafenib and ramucirumab) have been integrated into clinical practice^90^ (Figure 2).

The allele frequencies of variants that interact with antiangiogenic agents rs9582036, rs12505758, rs3025039, rs1049550 and rs2230054 in populations worldwide are shown in Table 957.

**Table 9.** Allele frequencies for clinically relevant germline polymorphisms VEGFR1 rs9582036, VEGFR2 rs12505758, VEGFA rs3025039, ANXA11 rs1049550 and CXCR2 rs2230054 in populations worldwide

| Gene | Polymorphism | Human Populations |  |  |
| --- | --- | --- | --- | --- |
|  |  | Latin American | Caucasian | Asian |
| VEGFR1 | rs9582036<br>(C-834A) | Colombia: 0.73 (A)<sup>a</sup>;<br>Mexico: 0.84 (A);<br>Peru: 0.94 (A);<br>Puerto Rico: 0.64 (A) | Spain: 0.69 (A);<br>British: 0.70 (A);<br>Finland: 0.78 (A);<br>Italy: 0.75 (A) | Han Chinese: 0.83 (A);<br>Bangladesh: 0.84 (A);<br>Japan: 0.85 (A);<br>Vietnam: 0.79 (A) |
| VEGFR2 | rs12505758<br>(2266+1166 A>G) | Colombia: 0.13 (C);<br>Mexico: 0.17 (C);<br>Peru: 0.31 (C);<br>Puerto Rico: 0.09 (C) | Spain: 0.12 (C);<br>British: 0.07 (C);<br>Finland: 0.10 (C);<br>Italy: 0.12 (C) | Han Chinese: 0.19 (C);<br>Bangladesh: 0.34 (C);<br>Japan: 0.27 (C);<br>Vietnam: 0.15 (C) |
| VEGFA | rs3025039<br>(C*237T) | Colombia: 0.13 (T);<br>Mexico: 0.30 (T);<br>Peru: 0.34 (T);<br>Puerto Rico: 0.18 (T) | Spain: 0.13 (T);<br>British: 0.07 (T);<br>Finland: 0.14 (T);<br>Italy: 0.12 (T) | Han Chinese: 0.18 (T);<br>Bangladesh: 0.12 (T);<br>Japan: 0.16 (T);<br>Vietnam: 0.16 (T) |
| ANXA11 | rs1049550<br>(C688T) | Colombia: 0.45 (A);<br>Mexico: 0.39 (A);<br>Peru: 0.64 (A);<br>Puerto Rico: 0.38 (A) | Spain: 0.47 (A);<br>British: 0.44 (A);<br>Finland: 0.54 (A);<br>Italy: 0.41 (A) | Han Chinese: 0.63 (A);<br>Bangladesh: 0.35 (A);<br>Japan: 0.65 (A);<br>Vietnam: 0.59 (A) |
| CXCR2 | rs2230054<br>(C786T) | Colombia: 0.45 (T);<br>Mexico: 0.45 (T);<br>Peru: 0.57 (T);<br>Puerto Rico: 0.54 (T) | Spain: 0.51 (T);<br>British: 0.47 (T);<br>Finland: 0.42 (T);<br>Italy: 0.50 (T) | Han Chinese: 0.36 (T);<br>Bangladesh: 0.49 (T);<br>Japan: 0.28 (T);<br>Vietnam: 0.39 (T) |

ANXA11, annexin A11; CXCR2, C-X-C motif chemokine receptor 2; VEGFA, vascular endothelial growth factor A; VEGFR1, vascular endothelial growth factor receptor 1; VEGFR2, vascular endothelial growth factor receptor 2; G, guanine; A, adenine; C, cytosine; T, thymine. <sup>a</sup> Frequency of minor allele.

#### Agents against epidermal growth factor receptor

Cetuximab and panitumumab are monoclonal antibodies (mAb) that block the action of EGF and may be employed in mCRC treatment^7, 92^. These drugs exert their action by binding to the extracellular domain of EGFR, with a greater affinity than the wild-type EGF, thereby blocking phosphorylation induced by EGFR ligands. Some variants are shown in Table 10^62, 63, 73, 100–108^.

**Table 10.**
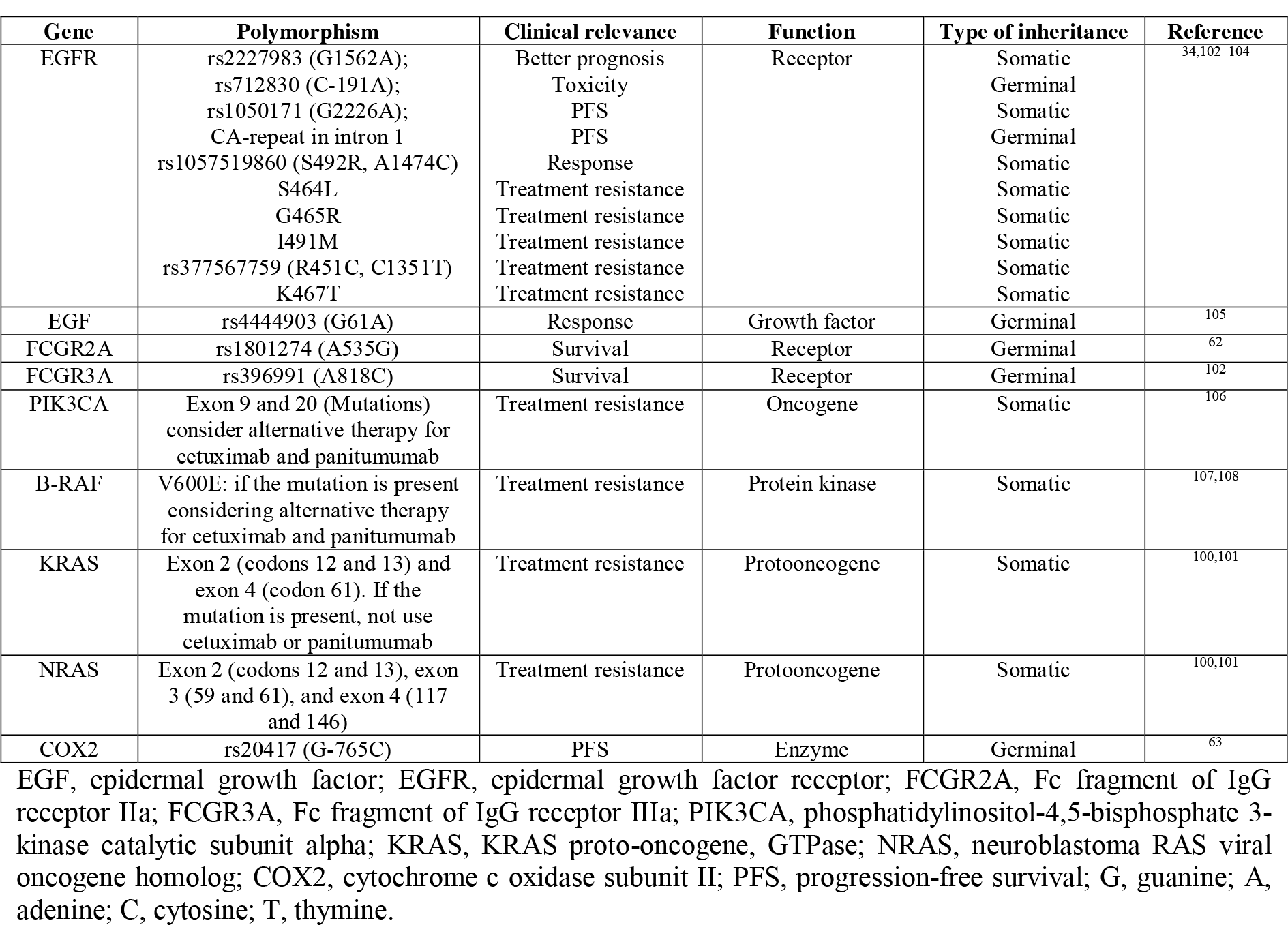
Biomarkers focused on gefitinib, erlotinib, imatinib, vemurafenib, cetuximab, and panitumumab drugs

The EGF/EGFR pathway plays an important role in cancer pathogenesis. EGF and EGFR are commonly overexpressed in CRC and they appear to be associated with poor prognosis and increased metastatic risk^109^. EGFR is a transmembrane glycoprotein that plays a main role in cell proliferation, migration and survival. EGFR R497K attenuates tyrosine kinase activation^110^, and EGF G61A increases its production when individuals have GG or GA genotypes^105^. The EGF/EGFR pathway is a predictive marker for cetuximab treatment in patients with locally advanced CRC^111^.

KRAS oncogene is a member of the human RAS family, which produces a self-inactivating guanosine triphosphate (GTP), binding signal transducer located on the inner surface of the cell membrane^112^. KRAS mutations may compromise the intrinsic GTPase activity, resulting in constitutively active KRAS protein that affects various signaling pathways^112^. The 45% of CRC cases has KRAS mutations and it has been shown that these mutations are predictive biomarkers of poor outcome in mCRC treated with cetuximab^77^. The anti-EGFR mAb therapy significantly improves both PFS and OS tumors without mutations in RAS. Therefore, mutations in KRAS predict resistance to mAb directed to EGFR with cetuximab and panitumumab^100^. NRAS codifies an isoform of RAS protein, involved primarily in the regulation of cell division^108^. Mutations in exon 2, 3 and 4 of NRAS, in addition to those in exon 2 of KRAS, must be detected before administration of a monoclonal anti-EGFR^100^. According to the NCCN, KRAS and NRAS are the only one predictive biomarkers approved in mCRC. Cetuximab and panitumumab are applied on patients with non-mutated RAS, and bevacizumab is applied on patients with mutated RAS^12, 113^.

Regarding Fc receptor range, modulating the immune response could be a further important mechanism to cetuximab sensitivity. The immune mechanism of antibody-dependent (ADCC) mediated cellular cytotoxicity through Fc receptors (Fc gamma R) made by immune cells, plays an important role in the effect of IgG1 antitumor antibodies^114, 115^. The most common polymorphisms in FCGR2A and FCGR3A are rs1801274 and rs396991, respectively^116, 117^.

The phosphatidylinositol 3-kinase (PI3K/AKT) pathway plays an essential role in cancer pathogenesis, being imperative the design of therapeutic inhibitors^118^. PIK3CA gene encodes the p110 catalytic subunit of PI3K alpha. PIK3CA mutations (associated with KRAS mutation and MSI) stimulate AKT pathway and promote cell growth in CRC^118^. PIK3CA mutations in exon 9 and 20 affects the helical and kinase domains of the protein, promoting a lack of effectiveness in drug treatments^106^.

BRAF protein is part of the RAS/MAPK signaling pathway, which regulates cell growth, proliferation, migration and apoptosis^119^. BRAF is a driver gene whose mutations are inversely associated with treatment response and are mutually exclusive with RAS mutations^108, 120^. Hence, BRAF V600E mutation correlates with worse prognosis^121^. Vemurafenib is a third-line therapeutic option in advanced mCRC with BRAF mutations^122^. Furthermore, it has been proposed that patients with KRAS and BRAF mutations could be eligible for mAb treatment against EGFR. Finally, BRAF must not present any mutation for a favorable treatment response when panitumumab or cetuximab are applied^107^.

COX is the limiting enzyme in the conversion of arachidonic acid into prostaglandins. COX2, encoded by prostaglandin endoperoxide synthase 2 (PTGS2), is involved in metastasis and chemotherapy resistance^123^. High levels of COX2 are associated with shorter OS in CRC. The C allele of COX2 G765C polymorphism has been associated with a significantly lower promoter activity^104, 124^.

The allele frequencies of rs2227983, rs4444903, rs1801274, rs396991 and rs20417 genetic variants in populations worldwide are shown in Table 11^57^.

**Table 11.** Allele frequencies for clinically relevant genetic variants EGFR rs2227983, EGF rs4444903, FGGR2A rs1801274, FCGR3A rs396991 and COX2 rs20417 in populations worldwide

| Gene | Polymorphism | Human Populations |  |  |
| --- | --- | --- | --- | --- |
|  |  | Latin American | Caucasian | Asian |
| EGFR | rs2227983 (G1562A) | Colombia: 0.35 (A)<sup>a</sup>;<br>Mexico: 0.31 (A);<br>Peru: 0.36 (A);<br>Puerto Rico: 0.30 (A) | Spain: 0.25 (A);<br>British: 0.24 (A);<br>Finland: 0.38 (A);<br>Italy: 0.25 (A) | Han Chinese: 0.46 (A);<br>Bangladesh: 0.34 (A);<br>Japan: 0.62 (A);<br>Vietnam: 0.53 (A) |
| EGF | rs4444903 (G61A) | Colombia: 0.51 (G);<br>Mexico: 0.62 (G);<br>Peru: 0.70 (G);<br>Puerto Rico: 0.52 (G) | Spain: 0.39 (G);<br>British: 0.41 (G);<br>Finland: 0.37 (G);<br>Italy: 0.38 (G) | Han Chinese: 0.70 (G);<br>Bangladesh: 0.64 (G);<br>Japan: 0.71 (G);<br>Vietnam: 0.70 (G) |
| FCGR2A | rs1801274 (A535G) | Colombia: 0.39 (G);<br>Mexico: 0.51 (G);<br>Peru: 0.47 (G);<br>Puerto Rico: 0.45 (G) | Spain: 0.53 (G);<br>British: 0.61 (G);<br>Finland: 0.54 (G);<br>Italy: 0.41 (G) | Han Chinese: 0.34 (G);<br>Bangladesh: 0.36 (G);<br>Japan: 0.19 (G);<br>Vietnam: 0.28 (G) |
| FCGR3A | rs396991 (A818C) | Colombia: 0.00 (C);<br>Mexico: 0.00 (C);<br>Peru: 0.00 (C);<br>Puerto Rico: 0.00 (C) | Spain: 0.00 (C);<br>British: 0.00 (C);<br>Finland: 0.00 (C);<br>Italy: 0.00 (C) | Han Chinese: 0.00 (C);<br>Bangladesh: 0.00 (C);<br>Japan: 0.00 (C);<br>Vietnam: 0.01 (C) |
| COX2 | rs20417 (G-765C) | Colombia: 0.22 (G);<br>Mexico: 0.21 (G);<br>Peru: 0.21 (G);<br>Puerto Rico: 0.21 (G) | Spain: 0.15 (G);<br>British: 0.14 (G);<br>Finland: 0.11 (G);<br>Italy: 0.19 (G) | Han Chinese: 0.05 (G);<br>Bangladesh: 0.17 (G);<br>Japan: 0.04 (G);<br>Vietnam: 0.02 (G) |

EGF, epidermal growth factor; EGFR, epidermal growth factor receptor; FCGR2A, Fc fragment of IgG receptor IIa; FCGR3A, Fc fragment of IgG receptor IIIa; COX2, cytochrome c oxidase subunit II; G, guanine; A, adenine; C, cytosine; T, thymine. <sup>a</sup> Frequency of minor allele.

#### Colorectal cancer immunogenomics

Recent advances in cancer immunology have highlighted the immunogenic nature of CRC and provided insights regarding the complex tumor-immune system interactions that drive immune evasion in CRC^125, 126^. One of the mechanisms that mediates tumor-associated immune escape is the activation of inhibitory co-receptors or immune checkpoints on the T lymphocyte surface by tumor cells through the expression of immunosuppressive molecules^125–127^.

Programmed cell death protein 1 (PD-1, CD279) is an inhibitory co-receptor expressed by exhausted tumor-infiltrating lymphocytes (TILs) present within the tumor microenvironment^127–129^. PD-1 engages with programmed-death ligands 1 (PD-L1, BT-H1, CD274) and 2 (PD-L2, B7-DC, CD273) which are expressed by CRC cells^130–133^. PD-1/PD-L1 interaction inhibits CD8^+^ T-cell activation, cytokine production, proliferation and cytotoxicity which suppresses the host immune response and allows CRC cells to proliferate and metastasize^127–129^.

Immune checkpoint inhibition has revolutionized cancer immunotherapy since it has proven to be very successful for treatment of melanoma and non-small cell lung cancer^134–136^. It has been shown that PD-1 blockade is a highly efficient therapeutic strategy against MSI-high and MMRd CRC tumors since these tumors display dense lymphocyte infiltrates due to their increased expression of immunogenic neo-antigens^137–139^. Moreover, these tumors exhibit higher PD-1 expression on TILs and PD-L1 expression than microsatellite stable tumors^139, 140^. Subsequently, the FDA approved pembrolizumab and nivolumab, two anti PD-1 antibodies, for treatment of metastatic MSI-high or MMRd solid tumors.

According to Tauriello *et al*., inhibition of the PD-1 and PD-L1 immune checkpoints provoked a limited response in quadruple-mutant mice. By contrast, his results strongly suggest that inhibition of TGFβ signaling could be promising as immunotherapy for patients with microsatellite stability and stroma-rich CRCs, enduring cytotoxic T-cell response against tumor cells that prevent metastasis^141–143^. The clinical implications of CRC immunogenomics continue to expand, and it will likely serve as a guide for next-generation immunotherapy strategies for improving outcomes for this disease (Figure 2).

### The Pan-Cancer Atlas: germline pathogenic variants

The Pan-Cancer Atlas provides a panoramic view of the oncogenic processes that contribute to human cancer. It reveals how genetic variants collaborate in cancer progression and explores the influence of mutations on cell signaling and immune cell composition, providing insight to prioritize the development of new immunotherapies^144^.

According to Huang *et al*., the Pan-Cancer Atlas analyzed 564 CRC samples and found several pathogenic germline variants in the APC, ATM, ATR, BARD1, BLM, BRCA1, BRCA2, BRIP1, CHEK2, COL7A1, FANCI, GJB2, MLH1, MSH2, MSH6, PALB2, POT1, RAD51D, RECQL4, RET, RHBDF2 and SDHA genes^145^. Additionally, Table 12 shows the allele frequencies of those 29 pathogenic germline variants according to The Exome Aggregation Consortium (ExAC)^146^.

**Table 12.**
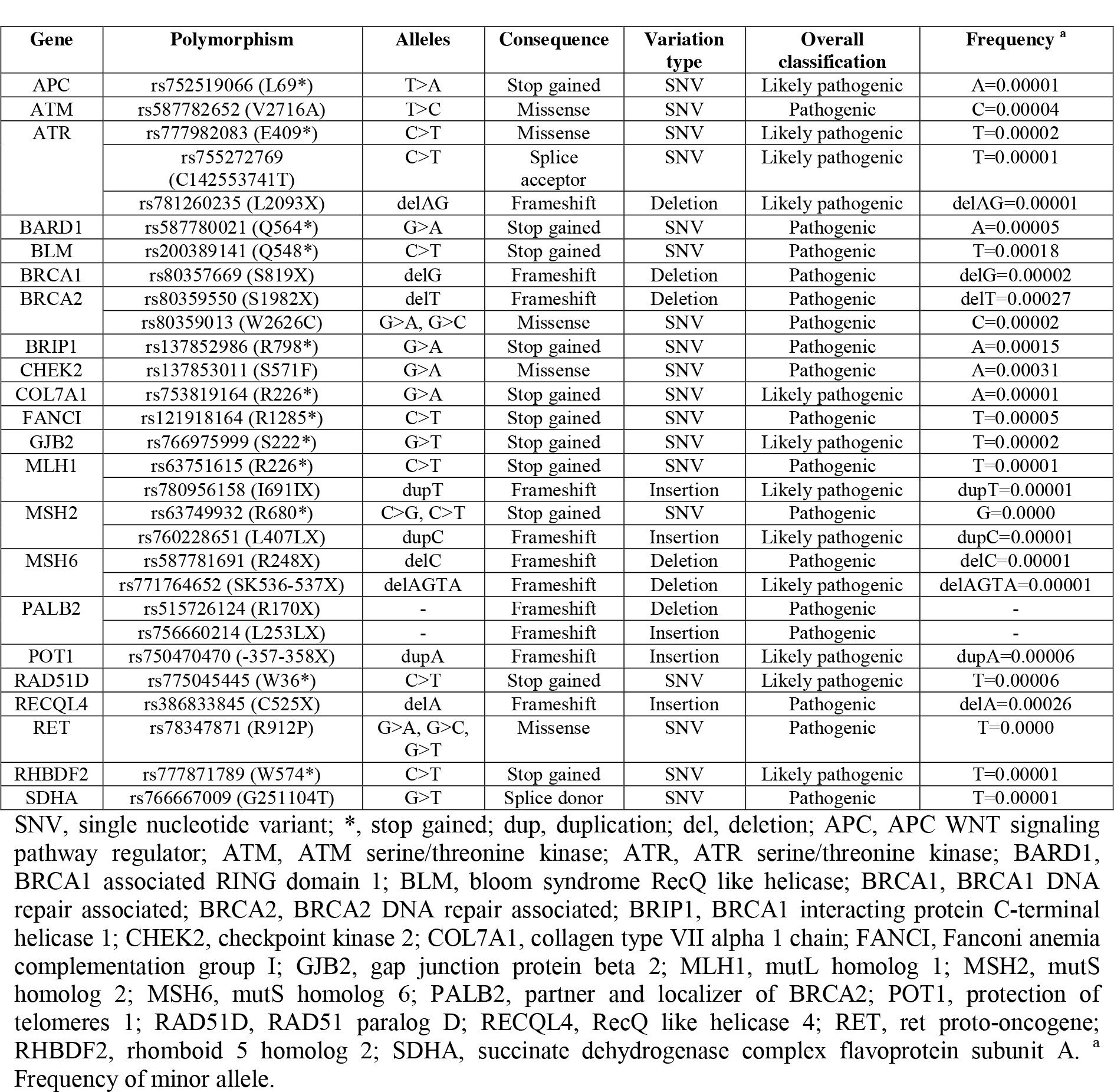
Pathogenic germline variants in CRC according to the Pan-Cancer Atlas and allele frequencies according to the Exome Aggregation Consortium

| Gene | Polymorphism | Alleles | Consequence | Variation type | Overall classification | Frequency <sup>a</sup> |
| --- | --- | --- | --- | --- | --- | --- |
| APC | rs752519066 (L69*) | T>A | Stop gained | SNV | Likely pathogenic | A=0.00001 |
| ATM | rs587782652 (V2716A) | T>C | Missense | SNV | Pathogenic | C=0.00004 |
| ATR | rs777982083 (E409*) | C>T | Missense | SNV | Likely pathogenic | T=0.00002 |
|  | rs755272769 (C142553741T) | C>T | Splice acceptor | SNV | Likely pathogenic | T=0.00001 |
|  | rs781260235 (L2093X) | delAG | Frameshift | Deletion | Likely pathogenic | delAG=0.00001 |
| BARD1 | rs587780021 (Q564*) | G>A | Stop gained | SNV | Pathogenic | A=0.00005 |
| BLM | rs200389141 (Q548*) | C>T | Stop gained | SNV | Pathogenic | T=0.00018 |
| BRCA1 | rs80357669 (S819X) | delG | Frameshift | Deletion | Pathogenic | delG=0.00002 |
| BRCA2 | rs80359550 (S1982X) | delT | Frameshift | Deletion | Pathogenic | delT=0.00027 |
|  | rs80359013 (W2626C) | G>A, G>C | Missense | SNV | Pathogenic | C=0.00002 |
| BRIP1 | rs137852986 (R798*) | G>A | Stop gained | SNV | Pathogenic | A=0.00015 |
| CHEK2 | rs137853011 (S571F) | G>A | Missense | SNV | Pathogenic | A=0.00031 |
| COL7A1 | rs753819164 (R226*) | G>A | Stop gained | SNV | Likely pathogenic | A=0.00001 |
| FANCI | rs121918164 (R1285*) | C>T | Stop gained | SNV | Pathogenic | T=0.00005 |
| GJB2 | rs766975999 (S222*) | G>T | Stop gained | SNV | Likely pathogenic | T=0.00002 |
| MLH1 | rs63751615 (R226*) | C>T | Stop gained | SNV | Pathogenic | T=0.00001 |
|  | rs780956158 (I691IX) | dupT | Frameshift | Insertion | Likely pathogenic | dupT=0.00001 |
| MSH2 | rs63749932 (R680*) | C>G, C>T | Stop gained | SNV | Pathogenic | G=0.0000 |
|  | rs760228651 (L407LX) | dupC | Frameshift | Insertion | Likely pathogenic | dupC=0.00001 |
| MSH6 | rs587781691 (R248X) | delC | Frameshift | Deletion | Pathogenic | delC=0.00001 |
|  | rs771764652 (SK536-537X) | delAGTA | Frameshift | Deletion | Likely pathogenic | delAGTA=0.00001 |
| PALB2 | rs515726124 (R170X) | - | Frameshift | Deletion | Pathogenic | - |
|  | rs756660214 (L253LX) | - | Frameshift | Insertion | Pathogenic | - |
| POT1 | rs750470470 (-357-358X) | dupA | Frameshift | Insertion | Likely pathogenic | dupA=0.00006 |
| RAD51D | rs775045445 (W36*) | C>T | Stop gained | SNV | Likely pathogenic | T=0.00006 |
| RECQL4 | rs386833845 (C525X) | delA | Frameshift | Insertion | Pathogenic | delA=0.00026 |
| RET | rs78347871 (R912P) | G>A, G>C, G>T | Missense | SNV | Pathogenic | T=0.0000 |
| RHBDF2 | rs777871789 (W574*) | C>T | Stop gained | SNV | Likely pathogenic | T=0.00001 |
| SDHA | rs766667009 (G251104T) | G>T | Splice donor | SNV | Pathogenic | T=0.00001 |

SNV, single nucleotide variant; \*, stop gained; dup, duplication; del, deletion; APC, APC WNT signaling pathway regulator; ATM, ATM serine/threonine kinase; ATR, ATR serine/threonine kinase; BARD1, BRCA1 associated RING domain 1; BLM, bloom syndrome RecQ like helicase; BRCA1, BRCA1 DNA repair associated; BRCA2, BRCA2 DNA repair associated; BRIP1, BRCA1 interacting protein C-terminal helicase 1; CHEK2, checkpoint kinase 2; COL7A1, collagen type VII alpha 1 chain; FANCI, Fanconi anemia complementation group I; GJB2, gap junction protein beta 2; MLH1, mutL homolog 1; MSH2, mutS homolog 2; MSH6, mutS homolog 6; PALB2, partner and localizer of BRCA2; POT1, protection of telomeres 1; RAD51D, RAD51 paralog D; RECQL4, RecQ like helicase 4; RET, ret proto-oncogene; RHBDF2, rhomboid 5 homolog 2; SDHA, succinate dehydrogenase complex flavoprotein subunit A. <sup>a</sup> Frequency of minor allele.

### Biomarker network in colorectal cancer

Figure 3 shows the proposed biomarker network in CRC. The protein-protein interaction (PPi) network with a highest confidence cutoff of 0.9 was created using String Database^147^. This network is made up of known and predicted interactions of driver genes, genes with pathogenic germline mutations according to the Pan-Cancer Atlas^33, 145^, genes with somatic mutations according to the CGI^34^, and druggable enzymes according to the Pharmacogenomics Knowledge Base (PharmGKB)^148, 149^.

**Figure 3.**
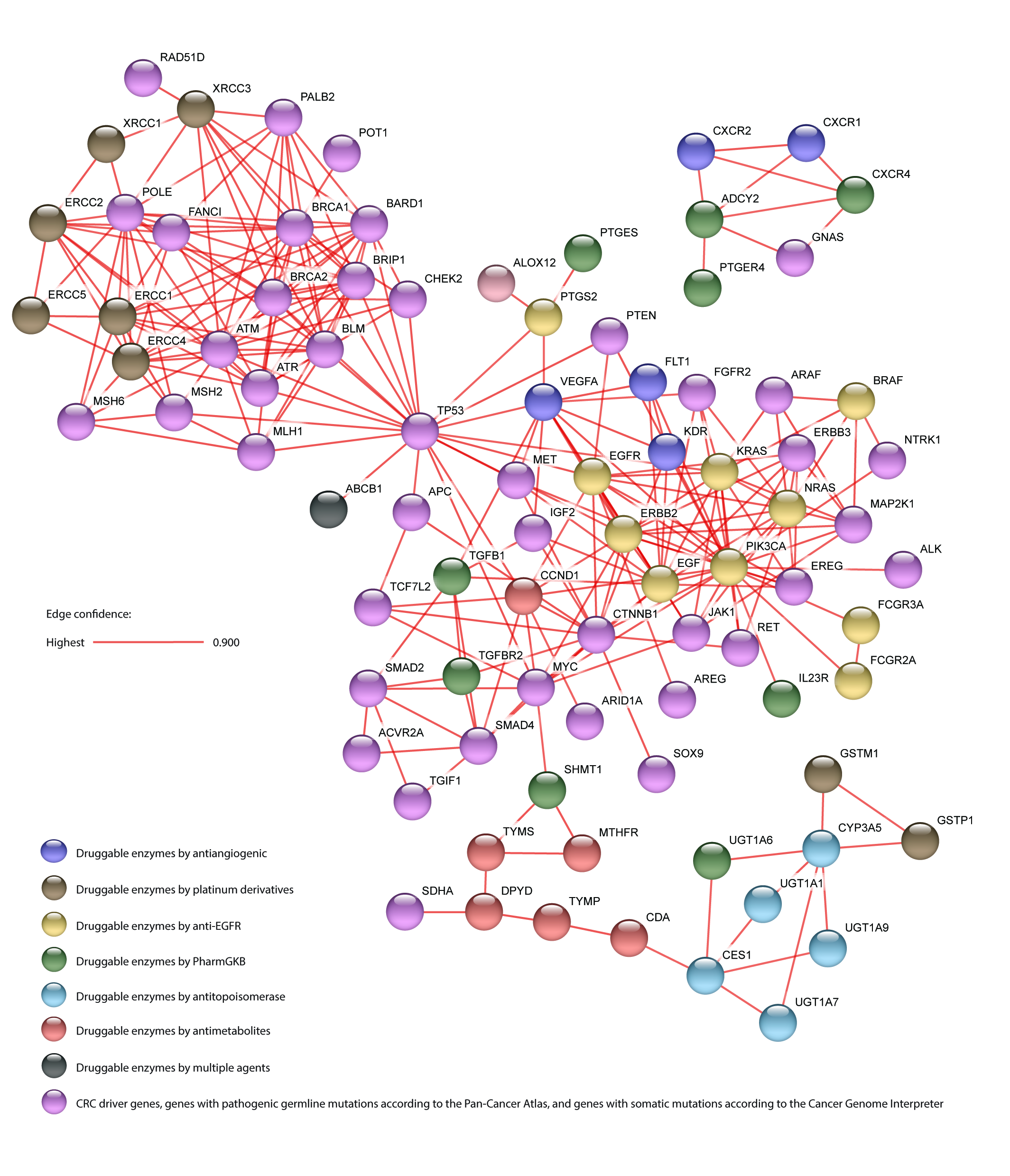
Biomarker network in CRC made up of driver genes, genes with pathogenic germline mutations, genes with somatic mutations, and druggable enzymes by antitopoisomerase, antimetabolite, platinum derivative and antiangiogenic drugs. The PPi network with a highest confidence cutoff of 0.9 was created using String Database.

The enrichment analysis of gene ontology (GO) terms related to biological processes and metabolic pathways were carried in the 87 genes of CRC biomarker network (Figure 3). The top biological processes with significant false discovery rate (FDR) <0.01 were DNA synthesis involved in DNA repair, strand displacement and response to drug. Meanwhile, the top metabolic pathways with FDR <0.01 were colorectal, endometrial and pancreatic cancer types^150^.

### Pharmacogenomics in clinical practice

In addition to the NCCN and ESMO guidelines^36–38^, the Canadian Pharmacogenomics Network for Drug Safety (CPNDS), the Royal Dutch Association for the Advancement of Pharmacy (DPWG) and the Clinical Pharmacogenetics Implementation Consortium (CPIC) have published precise guidelines for the application of pharmacogenomics in clinical practice^151–153^. All this information is published in the PharmGKB which is a comprehensive resource that curates knowledge about the impact of 80 clinical annotations on drug response^148^ (Supplementary Table 2).

In addition to the 1000 Genomes Project (Phase 3)^57^, there are 33 studies that have published the allele frequencies of mutations in several druggable enzymes in 34 different ethnic populations from Latin America (Supplementary Tables 3-10). Figure 4 is an innovative way to visualize and correlate the minor allele frequencies of 43 genes related to the different categories of drugs applied in CRC treatments in 8674 samples from 9 Latin American countries. This information will make it easier for clinical oncologists to make decisions regarding CRC treatments. For instance, the minor allele (G) of GSTP1 rs1695 can alter GST activity reducing detoxification capacity and leading to increased efficacy of platinum compounds^42, 43^. Thus, the Latin American countries whom best reduce the detoxification capacity and increase the efficacy of platinum compounds are Venezuela, Mexico and Peru due to their populations have a G allele frequency ≥ 0.50.

**Figure 4.**
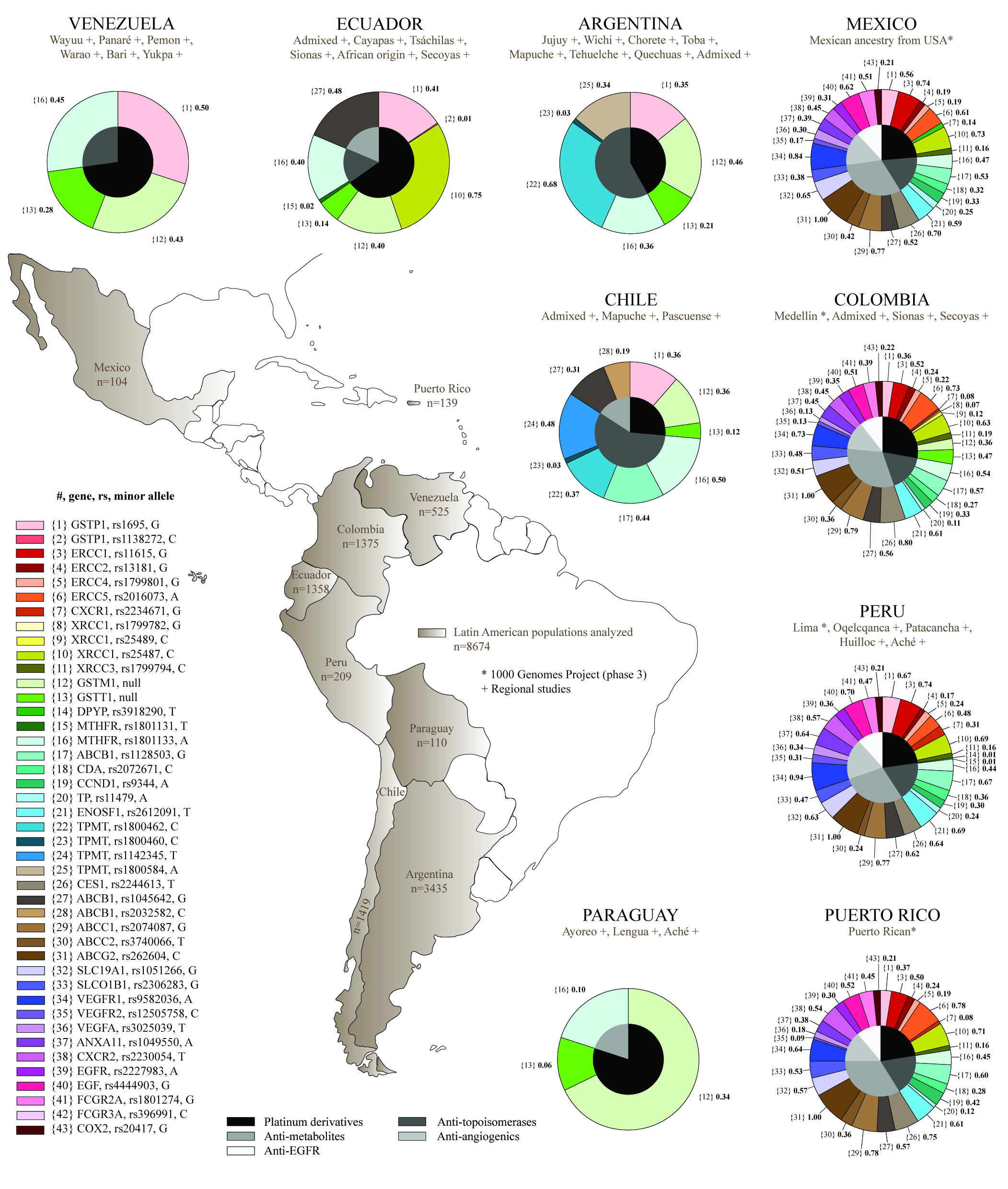
Minor allele frequencies of druggable enzymes studied in 8674 samples from Latin American populations, and its relation with the category of drugs applied in CRC treatments.

On the other hand, it is imperative to unify efforts at the governmental level to increase investment in pharmacogenomics fomenting precision medicine in clinical practice. The most relevant barriers to implement pharmacogenomics testing in clinical practice in Latin America are: 1) need for clear guidelines for the use of pharmacogenomics, 2) insufficient awareness of pharmacogenomics among clinicians, and 3) absence of a regulatory institution that facilitates the use of pharmacogenomics tests^154^. Overcoming the previously mentioned barriers, pharmacogenomics will make it possible to improve public health investment, patient safety and drug dosage in CRC treatments^155^.

## CONCLUSION

In the era of precision medicine, it is important to unify all current knowledge about the CRC biology to improve patient treatments. Large-scale projects worldwide have studied the multi-omics landscape of CRC by implementing the CMS classification and generating new therapeutic targets related to different populations worldwide. Developed countries might incorporate racial/ethnic minority populations in future cancer researches and clinical trials, and developing countries might invest to obtain a database of genomic profiles of their populations with the overall objective of linking pharmacogenomics in clinical practice.

## Acknowledgements

This research was supported by Centro de Investigación Genética y Genómica of Universidad UTE and the Latin American Society of Pharmacogenomics and Personalized Medicine.

## Author contributions

ALC and NWS conceived the subject and wrote the manuscript. ALC, GJK, DPIB, JGC, IAC and PGR did data curation and supplementary data. SG, CPyM, PEL, LAQ and JPC made a substantial contribution to the discussion of content. All authors reviewed and/or edited the article before submission.

## Data availability statement

All data generated or analysed during this study are included in this published article (and its Supplementary Information).

## Disclaimer

The contents of this publication are solely the responsibility of the authors and do not necessarily represent the official view of the National Institutes of Health.

